# Exposure to stressors and antimicrobials induces cell-autonomous ultrastructural heterogeneity of an intracellular bacterial pathogen

**DOI:** 10.1101/2020.09.14.297432

**Authors:** Marc Schulte, Michael Hensel, Katarzyna Miskiewicz

## Abstract

Despite being clonal, bacterial pathogens show a remarkable physiological heterogeneity during infection of host and within host cells. This diversity is reflected by distinct ultrastructural morphotypes in transmission electron microscopy (TEM). Gram-negative bacteria visualized at high resolution by TEM show a rather simple composition of cytoplasm with a centrally located nucleoid and large number of ribosomes. The cytoplasm is separated from the external environment by inner and outer membranes. In this study, we show that individual cells of *Salmonella enterica* serovar Typhimurium (STM) are ultrastructural divergent in standard culture conditions, as well as during their intracellular lifestyle in mammalian host cells. STM can basically be discriminated into two morphotypes based on the criterion of cytoplasmic density. We identified environmental conditions which affect cytoplasmic densities. Using chemical treatments and defined mutant strains, we were able to link the occurrence of an electron-dense type to oxidative stress and other noxes. Furthermore, ultrastructural analyses of STM during infection and fluorescence reporter analyses for cell viability were combined in a correlative light and electron microscopy approach. We provide evidence that two newly characterized ultrastructural types with lucent or dense cytoplasm represent viable cells. Moreover, the presence of electron-dense types is stress related and can be experimentally induced only when amino acids are available in the environment. This study sheds more light on diversities between individual bacteria in populations and possible physiological meanings like a stress response to explain the diversities discussed.

**Importance:** Bacterial pathogens show a remarkable resilience to adverse conditions during infection. Although being genetically identical, a clonal population may contain dead, dormant, slowly as well as rapidly proliferating cells. The physiological state of individual cells in a population may be analyzed by fluorescent probes or reporters. In contrast, reliable markers to interrogate single cells regarding viability, response to environmental cues, and exposure to antimicrobial compounds are sparse for ultrastructural approaches. For intracellular *Salmonella enterica* we observed distinct ultrastructural morphotypes. Using defined experimental conditions, these morphotypes were linked to reactions of bacteria to stressors or antimicrobials. The parameters defined here provide criteria for the interpretation of bacterial heterogeneity on the ultrastructural level.

## Introduction

Recent research on pathogenic bacteria revealed that cells react individually when exposed to adverse conditions due to differences in their physiological status (1–7). Classification of diversities and understanding of their biological function are crucial for designing new antimicrobials, which would overcome bacterial resistance without the risk of imposing selective pressure towards bacterial survival. Despite great progress in understanding the role of individual virulence factors for bacterial pathogenesis, the resilient bacterial survival in detrimental environmental conditions is still enigmatic (8–10). In the context of stress response to environmental cues, the formation of a dormant state with metabolic shifts, and change in cytoplasmic dynamics was postulated (11, 12). In general, bacterial cells appear structurally more complex than previously considered. For instance, the bacterial cytoplasm displays, in addition to high molecular crowding, unusual motility dynamics for differently-sized particles, properties of glass-phase, or transitions to solid-states (13, 14).

Transmission EM (TEM) is very potent to visualize the composition of bacterial envelopes, protein complexes, and has shed light on protein shell structures, revolutionizing the view on bacterial organelles (15, 16). Conventional TEM is broadly used as standard method to evaluate effects of bactericidal compounds (6, 17–21). In contrast, TEM was rarely applied to describe diversities of pathogenic bacteria at the single-cell level.

Bacterial cells visualized by EM demonstrate high ultrastructural variability, but the causes of this diversity are unknown (22–34). Distinct reactions to environmental stress can be a reason for such heterogeneity, as shown for aquatic microorganisms (35). However, direct links between physiological state, stress factors, and the bacterial ultrastructure have not been demonstrated. Identification of such links could delineate indicators of changes or circumstances critical for bacterial survival, to predict formation of persisters, to estimate sensitivities of populations, or to develop preventive strategies against bacterial infections. Hence, the investigation of different ultrastructural types and their frequencies may allow to predict prevailing environmental conditions, especially in the background of analyses of intracellular pathogens, or analyses of bacterial populations *in vivo* or free-living isolates.

The capability of *Salmonella enterica* serovar Typhimurium (STM) to survive harsh conditions in environments within and outside of mammalian hosts makes it a good model organism to reveal the basis of bacterial heterogeneity. STM is a foodborne pathogen, capable to pass the low pH barrier of the stomach. STM is exposed to host defense mechanisms and competing microbes in the host gastrointestinal tract. STM can invade epithelial cells, survive and proliferate in host cells, and can abuses phagocytes for spreading and replication within phagosomes. Within host cells, STM is enclosed by a specific vacuole (*Salmonella*-containing vacuole, SCV) and drives the formation of tubular membrane network (*Salmonella*-induced filaments, SIF), supporting its intracellular survival and progression (36–38). Moreover, STM is capable to survive and replicate within the host cell cytoplasm after escaping the SCV (39). Intracellular survival requires fast stress response and cellular reprogramming for protection and repair when facing strong bactericidal host defenses such as reactive oxygen species (ROS) (40–43).

In this study, we shed light on the consequences of environmental changes and exposure to stress factors for the bacterial ultrastructure. We demonstrate that heterogeneity, at the single-cell level, and ultrastructural homogeneity depend on the environment and can be experimentally induced. The approach presented here enables to link ultrastructural diversity with the physiological status of individual bacteria and environmental cues. For that, we combined classical microbiological assays with qualitative and quantitative TEM to study effects of induced oxidative stress and bactericidal conditions in STM wild type (WT) and the *sodAB* strain hypersensitive to ROS. Furthermore, we developed a strategy for fast correlative light and electron microscopy (CLEM) using high-voltage TEM of thick serial sections, and a fluorescent reporter for measuring bacterial metabolic activity. These results validate that different ultrastructural types are viable bacteria. The combination of ultrastructural studies at the single-cell level with fluorescent reporters is the next step towards an understanding of bacteria as individual organisms and their lifestyles.

## Results

Bacterial cells rapidly respond to changing environments in order to adapt and to survive. We asked if different physiological states of bacteria are reflected by ultrastructural features. We reasoned that the bacterial response to different environments and to stressor or antimicrobial agents, may result in cells differing in their nanostructure.

### *Ultrastructural diversity of* Salmonella enterica *cells in culture*

First, we applied conventional TEM with and without post-contrasting with heavy metals. STM WT was grown in Phosphate-Carbon-Nitrogen (PCN, (44)) minimal medium at pH 7.4 for 3.5 h for culture at reduced growth rate of 0.98, compared to 1.33 in rich medium LB ((41, 45). TEM analysis revealed that bacteria were very similar in ultrastructure, with a clearly visible outer and inner membrane separated by the periplasmic space (Figure 1A). The distance between the outer and inner membranes was 20 ± 5 nm (mean ± SD). The center of cells, outlined by the inner membrane, was slightly electron dense with a difference of 135 ± 16 in mean gray values (MGV) to the background, referred further as electron density. That region contained cytoplasm with protein complexes like ribosomes, visible as denser particles, which were distributed homogenously within the cell. Electron lucent and ribosome-free regions consisted of up to 16% of total cytoplasmic area and occupied not larger than 28 nm^2^ of area of clearly visible nucleoids. Hence, bacteria grown in the minimal PCN medium at physiological pH formed uniform populations of similar ultrastructure.

**Figure 1:**
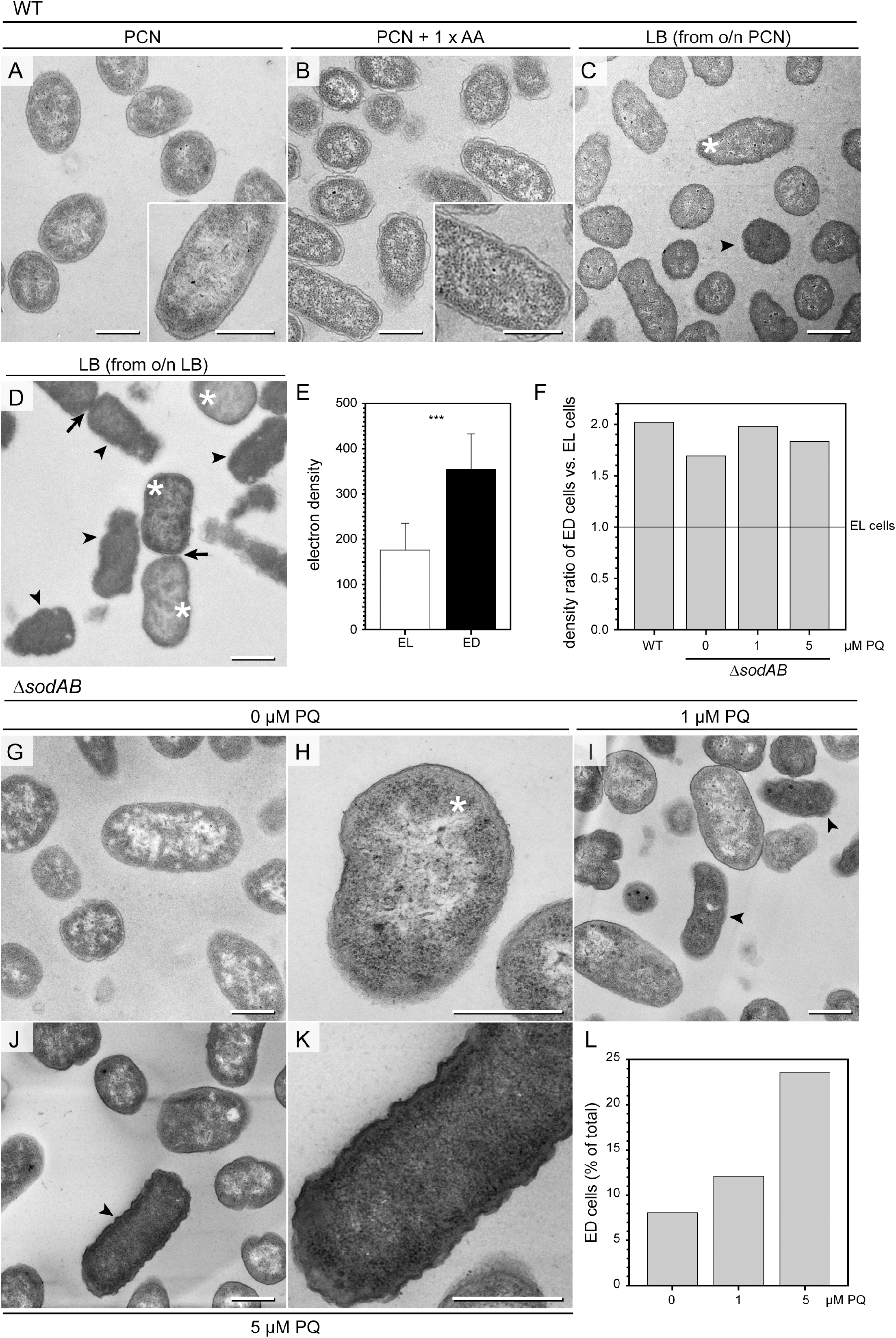
Environmental stress affects STM ultrastructure. **A-D**) Energy-filtered TEM (EF-TEM) micrographs (120 keV) showing electron-lucent STM WT after growth in PCN medium (**A**), PCN medium supplemented with AA (**B**), and electron-lucent (asterisks) and electron-dense (arrowheads) STM WT in LB medium subcultured for 3.5 h (**C, D**). Arrows indicate cells undergoing division. **E**) Comparison of STM WT densities in LB medium shown as mean ± SD of different values between bacterial cytoplasm mean grey values (MGV) and background MGV (see Fig. S1A, B, pooled data). **F**) Comparison of density ratio: electron-dense (arrowheads in **C**, **D**, **I**, **J**) vs. electron-lucent (asterisks in **C**, **D**, **H**) STM WT and Δ*sodAB* cultured in LB medium without or after PQ treatment. Numbers of cells quantified: 30, 22, 32, and 33, for STM WT, Δ*sodAB* 0 μM PQ, Δ*sodAB* 1 μM PQ, and Δ*sodAB* 5 μM PQ, respectively. **G-K**) TEM micrographs showing STM Δ*sodAB* without (**G**, **H**), or after treatment with PQ at 1 μM (**I**), or 5 μM (**J**, **K**). **L**) Comparison of the relative numbers of STM Δ*sodAB* ED. Numbers of cells quantified: 211, 157, and 135, for 0 μM, 1 μM, and 5 μM PQ, respectively. Scale bars, 500 nm. Statistical analysis was accomplished by Student’s *t*-test and significance levels are indicated as follows: *, p < 0.05; **, p < 0.01; ***, p < 0.001; n.s., not significant.

In addition, we analyzed the ultrastructure of STM cells cultured in PCN pH 7.4 medium supplemented with an amino acid (AA) mix (44) to increase the bacterial growth rate (Figure 1B). STM grown in PCN medium containing AA resembled bacteria grown in PCN medium without AA supplementation and formed a uniform bacterial population (Figure 1).

This was in contrast to bacteria grown in lysogeny broth (LB) as standard rich medium. If bacteria grown overnight in PCN medium were subcultured for 3.5 h in LB medium, cells with highly electron-dense cytoplasm were found, referred to as ‘electron-dense’ (ED) cells (arrowhead in Figure 1C), in addition to cells similar to these grown in PCN medium, referred to as ‘electron-lucent’ (EL) cells (asterisk in Figure 1C). If bacteria grown overnight in LB medium were subcultured for 3.5 h in LB medium, few EL cells (asterisk in Figure 1D) and numerous ED cells (arrowheads in Figure 1D) were detected. The averaged cytoplasmic electron density was 176 ± 60 MGV and 354 ± 78 MGV for EL and ED cells, respectively (Figure 1E). In addition, we quantified differences in the MGV of the bacterial cytoplasm and the background per image, revealing higher values of electron densities for ED cells independently in every frame (Figure S1A). Their averaged value (Figure S1B) was very similar to the mean of pooled data, with a significant difference of electron density between EL and ED cells (Figure 1E and S1B-C). Moreover, nanostructures as well as the periplasm were indistinguishable in ED cells, contrary to EL cell ultrastructure. Both cell types were visible during cell division as an evidence of their high viability (Figure 1D, arrows). In addition, in stationary LB cultures (Figure S1EG), we found some dying cells, with partially or completely loss of inner membranes, signs of molecular condensation (dark spots, arrows in Figure S1E), and/or lysis (the arrowhead in Figure S1G). These profiles also were frequent in STM WT of stationary PCN cultures, however, independently of medium pH or nutritional supplementations (Fig S1IKM). In 3.5 h subcultures of corresponding media, dying profiles were only sporadically found.

Hence depending on growth conditions, bacterial populations consist of either ultrastructural similar cells, or cells divergent in cytoplasmic electron densities.

### Controlled induction of STM ultrastructural diversity

In order to find a correlation between ultrastructural types and environmental stress factors, we deployed a STM Δ*sodAB* strain deficient in both cytoplasmic superoxide dismutases SodA and SodB. STM Δ*sodAB* is especially sensitive to oxidative stress and turned out to be fragile, being able to grow in LB rich medium, but not in PCN minimal medium (41). Populations of STM Δ*sodAB* showed ED and EL cells with ultrastructural features as STM WT (Figure S1F-G and 1F), however the EL cell type was dominant (Figure 1G-H, star). Next, we induced oxidative stress by adding methyl viologen (‘paraquat’, PQ), a redox-active compound producing superoxide. In the presence of PQ, STM Δ*sodAB* expose to this toxic radical was prolonged. Treatment for 1 h with PQ at low concentrations of 1 or 5 μM did not affect STM Δ*sodAB* growth on agar plates (similar number colony forming units, CFU). TEM revealed that these treatments caused increase in frequency of ED cells (Figure 1I-L, arrowheads) in a dose-dependent manner, supporting its specificity to PQ treatment (Figure 1L). These results suggest that ED cells represent a type responding to cellular stress induced i.e. by ROS.

To further scrutinize the link between ultrastructure and cellular stress, we analyzed STM Δ*sodAB* for abnormalities. STM Δ*sodAB* showed abnormal colony growth on agar plates when compared to STM WT, forming evidently smaller colonies at comparable number (Figure 2AB). Slower colony growth could be a result of cell division defects, high level of cell death and/or just slower growth. As revealed by TEM, cells of STM Δ*sodAB* were rod-shaped and of similar size to STM WT without any signs of higher cellular death (Figure S1D-G). In addition, we assessed the membrane integrity using propidium iodide (PI) (Figure 2C-N). PI is a red-fluorescent dye binding to DNA, which is not membrane-permeable, thus DNA staining reports membrane damages (46). STM Δ*sodAB* showed up to 5% of PI-positive cells at average without treatment (Figure 2C and S2), and even less PI-positive cells directly after incubation with 1 or 5 μM PQ (Figure 2DFH and S2A). We performed PI staining again 12 h after PQ treatment and found high numbers of PI-positive cells in a group treated with 5 μM PQ, suggesting membrane stress and initiation of progressive membrane injury (Figure 2EGH). Furthermore, ultrastructural analysis of STM Δ*sodAB* revealed high number of cells with membrane invaginations. These were often asymmetrical, and single or multiple events occurred, which were differently located also including cell poles, therefore representing abnormal cell envelopes. These features were also present in Δ*sodAB* without treatment, but increased after PQ treatment, suggesting oxidative stress as cause of this defect (Figure 2I-M, arrowheads). After treatment with 5 μM PQ, high-resolution analysis revealed cell profiles with damaged inner and outer membranes, manifested by a loss of integrity and a leakage of cytoplasmic content, (Figure 2NO), in a line with PI analysis. Foci of lysis were also present after PQ treatment (asterisks in Figure 2LN). Hence, PQ treatment of STM Δ*sodAB* enhanced cellular stress and could be toxic to cells.

**Figure 2:**
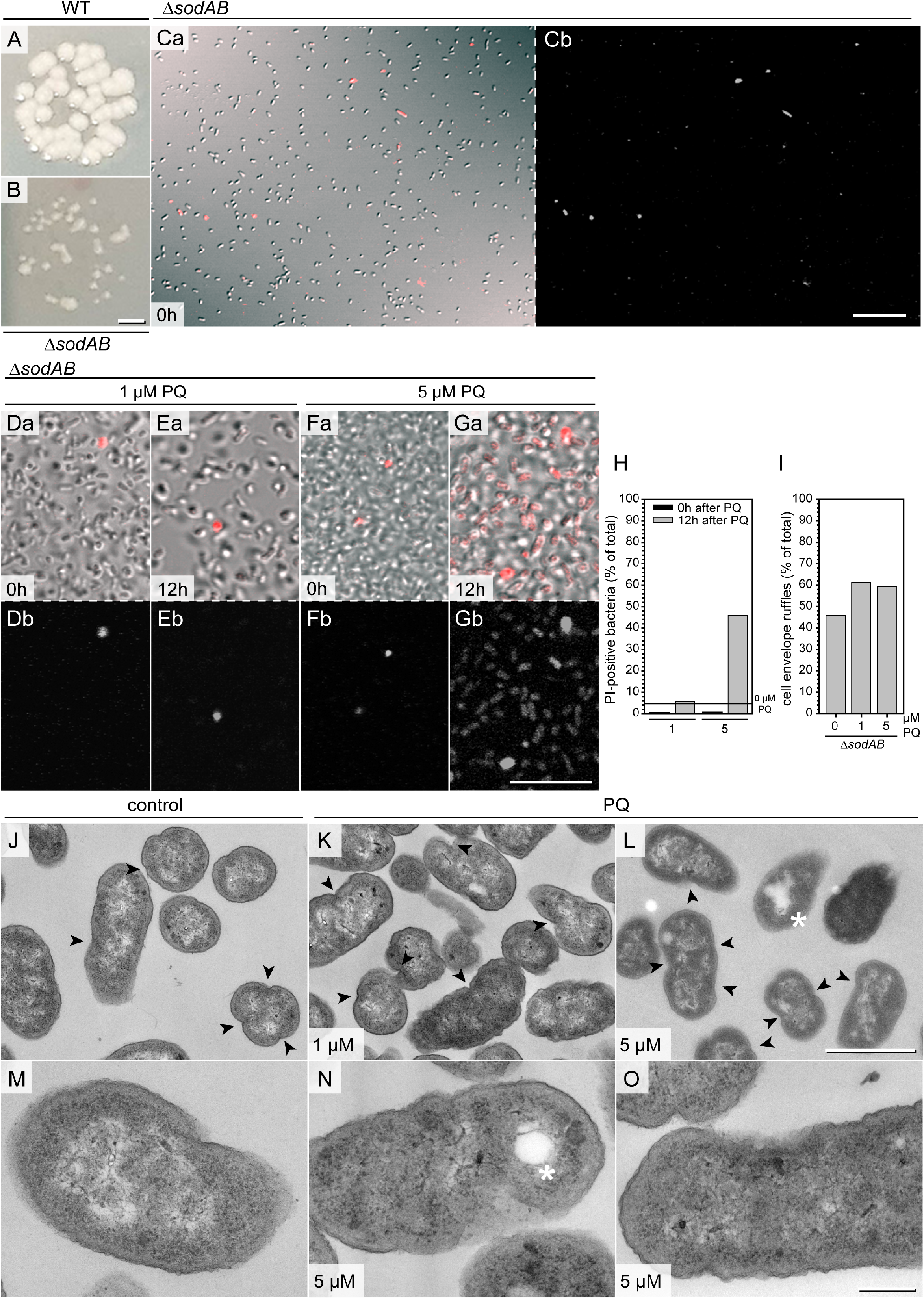
STM Δ*sodAB* has growth defects and exhibits membrane abnormalities. **A, B**) Growth of STM WT (**A**) and STM Δ*sodAB* (**B**) on agar plates. **C-H**) PI staining of STM Δ*sodAB* cultured for 3.5 h in LB medium without treatment (**C**), after treatment with 1 μM PQ (**D, E**), or 5 μM PQ (**F, G**) prior to TEM. **D** and **F** show bacteria with PI addition at time 0 h after PQ treatment, while **E** and **G** show cells with PI addition 12 h after PQ treatment. **H**) Quantification of PI-positive STM Δ*sodAB* cells. The line in **H** represents the level of PI-positive cells in untreated sample (related to Fig. S2). Number or quantified cells: 13,468, 1,565, 6,162, and 2,133, for 1 μM PQ at 0 h, 1 μM PQ at 12 h, 5 μM PQ at 0 h, and 5 μM PQ at 12 h, respectively. **I**) Relative numbers of STM Δ*sodAB* with cell envelope invaginations (arrowheads in **J-L**) of untreated control (**J, M**), or treated with 1 μM (**K**) or 5 μM PQ (**L, N, O**). **J-O**) TEM analysis by 120 keV EF-TEM of STM Δs*odAB* shown in **C, D, F**, fixed at 0 h post PQ treatment. Numbers of cells quantified: 113, 93, and 98, for 0 μM, 1 μM, and 5 μM PQ, respectively. **N**, **O**) STM Δ*sodAB* treated with 5 μM PQ shows membrane ruptures and lysis spot (asterisks). Scale bars, 10 μm (**C-G**), 1 μm (**J-L**), 200 nm (**M-O**).

### Induction of ultrastructural heterogeneity of STM WT in PCN medium

We confirmed that PQ concentrations higher than 50 μM were potent to induce membrane stress in STM WT in a similar fashion as in Δ*sodAB*, significantly raising the number of PI-positive cells and PI fluorescence intensity (Figure S2B). Exposure to 50 μM PQ achieved almost 70% PI-positive cells directly after treatment (Figure S2A). As pathogenic bacterium, STM possesses multiple stress response systems that activate repair mechanisms to protect from oxidative stress, thus increasing the chance to survive PQ treatment (47). Therefore, we defined a toxic PQ concentration which reduced the number of viable STM, and analyzed the ultrastructure of STM WT after PQ treatment in various growth conditions (Figure S3). PCN medium at pH 5.8 was also used to mimic the acidic phagosomal lumen of macrophages, where superoxide is protonated, capable to pass bacterial membranes but less spontaneously dismutates into hydrogen peroxide (2, 48). STM WT grown o/n in PCN medium at pH 7.4 was inoculated in the same medium and grown further for 3.5 h. PQ treatments were always performed in PCN medium with reduced concentration of inorganic phosphate (P_i_) of 0.4 mM, since P_i_ could compete with PQ during transport through bacterial membranes (49). A shift from the growth medium to a medium of acidic pH had lesser impact on colony growth, in comparison to bacteria shifted to medium of 7.4 pH, 96% of STM WT survived shift to pH 5.8, compared to shift to pH 7.4. After treatment with 100 μM PQ, significant drop of survival to 20% or less of controls was observed at both pH values. Treatments with higher PQ concentrations such as 500 μM or 1 mM, further reduced survival of STM WT, however, remained above 10% of controls (Figure S3W). TEM analysis of PQ-treated samples and controls revealed that in all conditions bacterial cells were uniform in ultrastructural appearance when cultured in PCN at pH 7.4 or pH 5.8 without AA supplementation (Figure 3AB and S3A-F). After PQ treatment, the cytoplasm was denser but ultrastructures like inner membranes, ribosomes or DNA were easily distinguishable. We did not find ED cells and we also did not find many cells with profiles of ultrastructural abnormalities (Figure S3A-L). It is possible that in PCN medium with limited nutrients, bacteria were not capable to switch to an emergency mode after PQ treatment, what would explain absence of ED cells and poor growth on agar plates.

**Figure 3:**
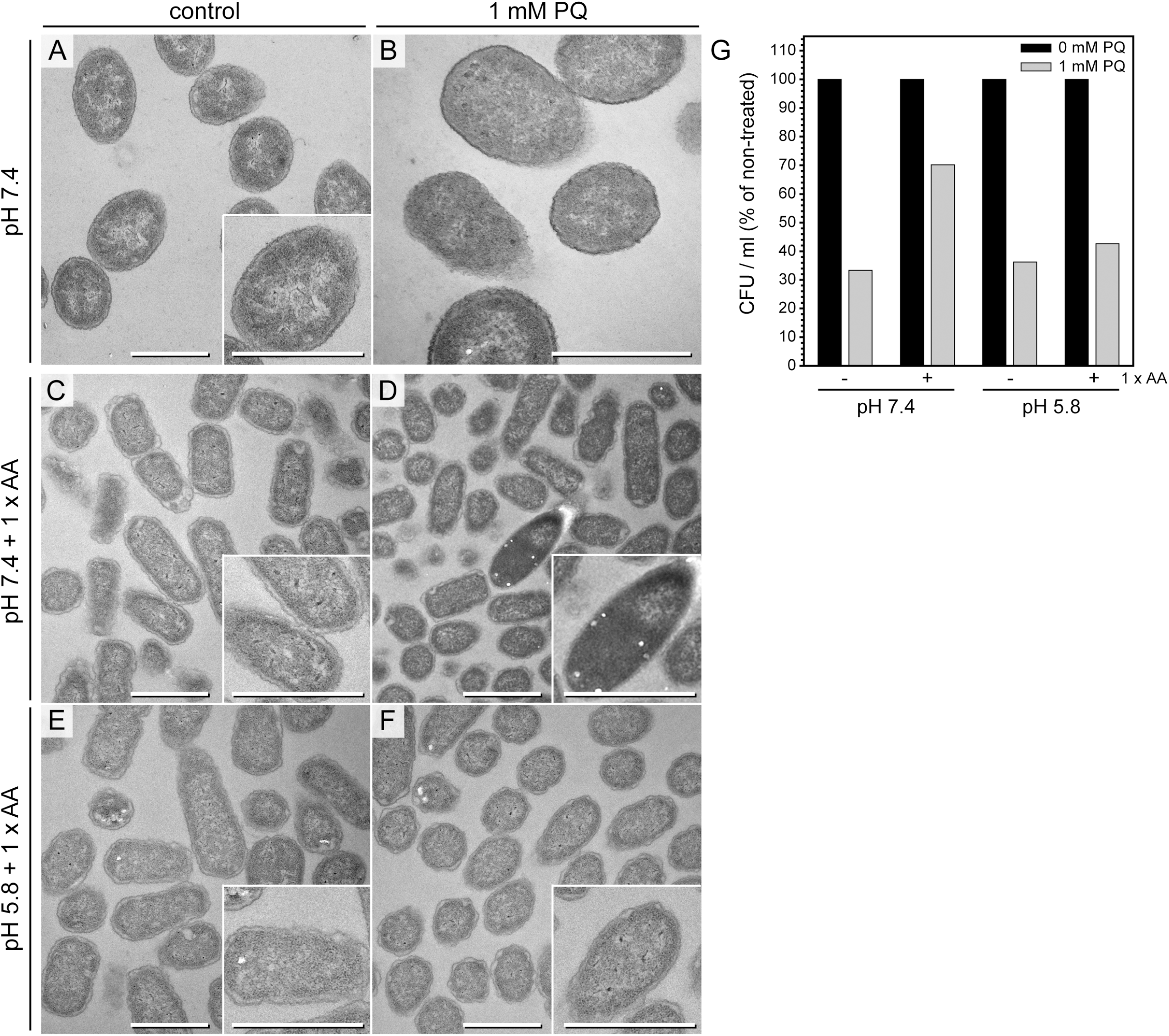
STM WT shows ED cells after PQ treatment in PCN pH 7.4 medium supplemented with AA. **A**, **B**) TEM micrograph of STM WT cultured in PCN medium, pH 7.4 for 3.5 h and shifted to fresh PCN medium, pH 7.4 for incubation without (control **A**) or with 1 mM PQ (B). **C**-**F**) Electron micrograph of STM WT cultured in PCN medium pH 7.4 (**C** and **D**), or pH 5.8 (**E** and **F**) supplemented with AA for 3.5 h and shifted to the same fresh PCN medium for incubation without (control **C** and **E**), or with 1 mM PQ (**D** and **F**). Scale bars, 1 μm. **G**) CFU counts obtained for STM WT without or with addition of 1 mM PQ. STM was subcultured for 3.5 h in PCN with or without AA supplementation, at pH 7.4 or pH 5.8.

To test this hypothesis, we performed the same experiments in PCN medium supplemented with AA (Figure 3C-F). STM survival after 1 mM PQ treatment was only higher when cultured in PCN medium pH 7.4 supplemented with AA, in contrast to STM grown without AA, or in PCN medium with a pH of 5.8 (Figure 3G). This was in line with the presence or lack of bacteria with ED type. ED cells emerged only in PCN pH 7.4 medium when supplemented with AA (Figure 3D). Treatments with lower PQ concentrations did not affect STM ultrastructure in PCN medium supplemented with AA (Figure S3M-O). For comparison, we also investigated the impact of other stress conditions on STM ultrastructure (Figure S3P-V). ED type was not observed after an osmotic shock or heat shock. It occurred after acid shock (shift to pH 3.0) of STM subcultured in PCN, pH 7.4 with AA supplementation, but not after subculture in PCN, pH 5.8. We observed other ultrastructural features, which were shock-specific to respective shock conditions and never observed in bacteria of control cultures. We compared presence of bacteria with shrinkage and/or lysis features since such profiles were observed in normal growth conditions (Figure S3Y). Cells with shrinkage and/or lysis features dominated the population after hyper osmotic stress in presence of 600 mM NaCl (shrinkage: 82.8-fold increase compared to untreated and 74.6% of total cells; lysis: 18.7-fold increase compared to untreated, 16.9% of total) (Figure S3U). Obvious increase of STM with signs of shrinkage and/or lysis was also observed after pH shock (pH 3.0) and was more pronounced when cells were subcultured at neutral pH. Simultaneous treatment with 1 mM PQ during pH shock resulted in comparable frequencies, suggesting only minor or no impact of PQ on causing shrinkage or lysis. This was in line with a low frequency of signs of shrinkage and/or lysis (< 10%) after PQ treatment in all other tested conditions (Figure S3Y) Hence, occurrence of ED type is induced by environmental stress and requires presence of AA.

### Ultrastructural diversity of intracellular STM

STM is able to replicate within eukaryotic cells where it encounters host cell defense mechanisms, as well as harsh phagosomal environments and nutritional limitations (40, 50). To correlate the ultrastructural features to intracellular phenotypes, we examined the ultrastructure of STM in HeLa cells at 8 h or 16 h post infection (p.i.). At both time points, host cells were either intact, with or without intracellular STM, or dying and ruptured as result of bacterial hyper-replication. Within healthy host cells, we found EL STM WT as well as mixed populations with EL and ED cells similarly to STM in LB medium (Figure 4). Both types were located within SCVs and showed signs of cell division. These data confirmed that ultrastructural EL and ED types are natural morphotypes of STM.

**Figure 4:**
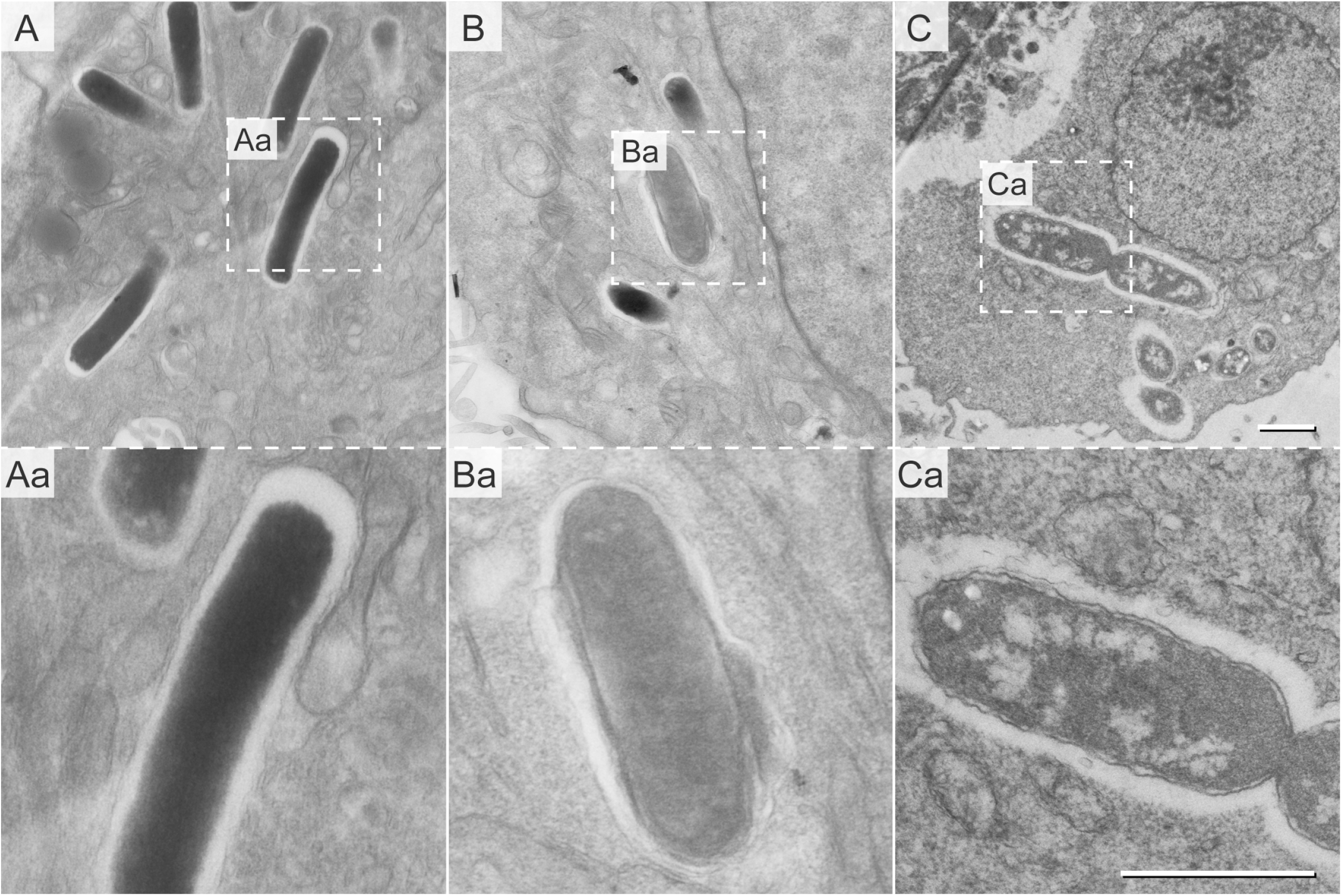
Ultrastructural diversity of intracellular STM WT. STM WT was subcultured in LB for 3.5 h and used to infect HeLa cells. Infected cells were fixed 16 h p.i. and analyzed by EF-TEM (120 keV). Micrographs show electron-dense (**A**) and electron-lucent STM WT (**B**, **C**). Dividing STM WT cells shown in **C**. Dashed boxes indicate areas enlarged in **a**. Scale bars, 1 μm.

### Metabolic activity of intracellular STM

In order to test vitality and biosynthetic capability of distinct morphotypes of STM, we used an episomal encoded dual-color fluorescence reporter (Figure S4A) (51). Bacteria harboring the reporter constitutively express *gfp*. To report viability and biosynthetic capacity, we monitored *dsred* expression regulated by the anhydrotetracycline (AHT)-inducible *tetA* promoter (52). We considered cells as biosynthetic active when DsRed was detected after AHT induction.

First, we tested reporter functionality by flow cytometry (Figure S4BCD). STM WT harboring the reporter was subcultured to late-logarithmic growth phase in LB medium. AHT was added after 3 h of growth, with or without chloramphenicol (Cm) to inhibit protein biosynthesis DsRed-positive cells were already detected 0.5-1 h after AHT induction. DsRed fluorescence intensity increased rapidly at 1.5-3 h post induction. Without AHT induction, or AHT induction in presence of Cm, no DsRed-positive cells were detected, verifying that inducible expression of *dsred* can be used as a marker for biosynthetic capability (Figure S4CD). We were also able to visualize metabolically active STM by live-cell fluorescence microscopy (FM) of infected host cells, supporting that the reporter can be used to study the metabolic status of intracellular STM (Figure S4E). We investigated infected HeLa LAMP1-GFP cells 8 h or 16 h p.i. (Figure 5). The intracellular population was heterogeneous based on protein synthesis and divergent when compared between infected host cells consisting of either only metabolically active or mixed metabolically active and inactive STM.

**Figure 5:**
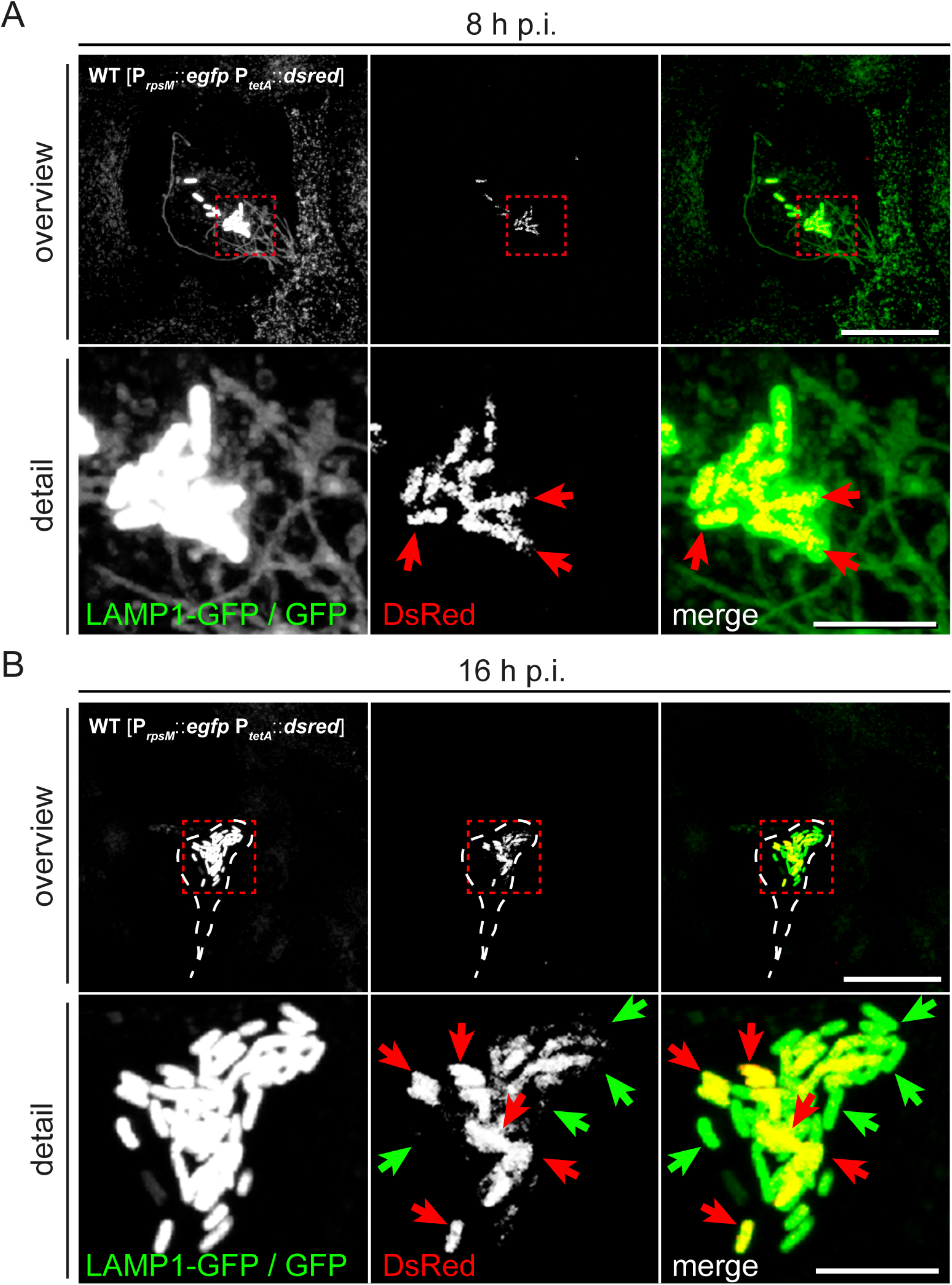
Hyper-replicating intracellular STM WT forms subpopulations of metabolically active and inactive bacteria. STM WT harboring dual-color vitality reporter was visualized inside HeLa LAMP1-GFP cells 8 h and 16 h p.i. *Gfp* was constitutively expressed in STM WT, while *dsred* expression was induced by addition of AHT 2 h prior to imaging. **A**) DsRed is visible in all intracellular STM WT associated with SIF formation 8 h p.i. (yellow cells in merge, red arrows). **B**) At 16 h p.i., hyper-replicating intracellular STM WT either lack DsRed (inactive cell indicated by green arrows) or are DsRed-positive (active cells indicated by red arrows). Scale bars, 20 and 5 μm in overview and detail, respectively.

Previous TEM analysis showed that host cell viability decreased when containing high burden of intracellular bacteria. Using LAMP1-GFP as marker we could assess *Salmonella*-induced endosomal remodeling (Figure 5A). We also observed lack of these compartments (Figure 5B) possibly due to activation of death processes in the host or due to their rupture by escaping STM into the host cell cytoplasm. Presence of individual metabolically inactive STM within a population of metabolically active STM raised the possibility that inactive STM are viable. Bacteria may form persisters with ceased growth, highly reduced metabolism, and ability to return to normal growth after release from stressful conditions. To scrutinize the bacterial conditions in host cells further, we applied correlative light and electron microscopy (CLEM).

### CLEM with the dual-color reporter strains

We modified a previous CLEM approach to accelerate data collection and further applied deconvolution of FM data (Figure S5) (53). We observed intracellular STM populations consisting of ED and EL cell types (72.7% ED cell type, 27.2% EL cell type), which were also visualized during cell division (Figure 6A, Ac). Highly electron-lucent single bacteria were found just once per ROI showing clearly visible outer membrane, outlined periplasm, and produced DsRed at high level (Figure 6Ad, Ba-b). However, there was no strict correlation of metabolically active, less active, or non-active STM with any of electron density-based morphotypes. We found both, ED and EL types strongly (34.4% of ED and 41.7% of EL) or slightly (18.8% of ED and 16.7% of EL) marked for DsRed expression. In addition, 46.9% of ED and 41.7% of EL cells showed metabolic inactivity. In ROIs with high numbers of STM, we detected clear differences in bacterial size, with an area of 1.76 ± 0.25 μm^2^ (wide) or 0.96 ± 0.18 μm^2^ (thin), which were only partially correlated with the electron density type (Figure 6A). Interestingly, size-based classes were rather grouped in the host cell with ‘wide’ STM located more centrally and ‘thin’ STM located on the cell peripheries (Figure 6A). Thin STM were highly metabolically active, while ED wide cells had no or minor expression of DsRed, localized to small patches. Dividing STM of both, ED and EL types, were wide and less metabolically active. Some bacteria were devoid of SCV and located in the host cell cytoplasm (Figure 6Ac). Complete SCV or SIF were not present, in line with lack of LAMP1-GFP signal. Contrary, lysosomes, endosomes and autophagic structures were present in a region with many thin and metabolically active STM, however, were not visible within degradative organelles.

**Figure 6:**
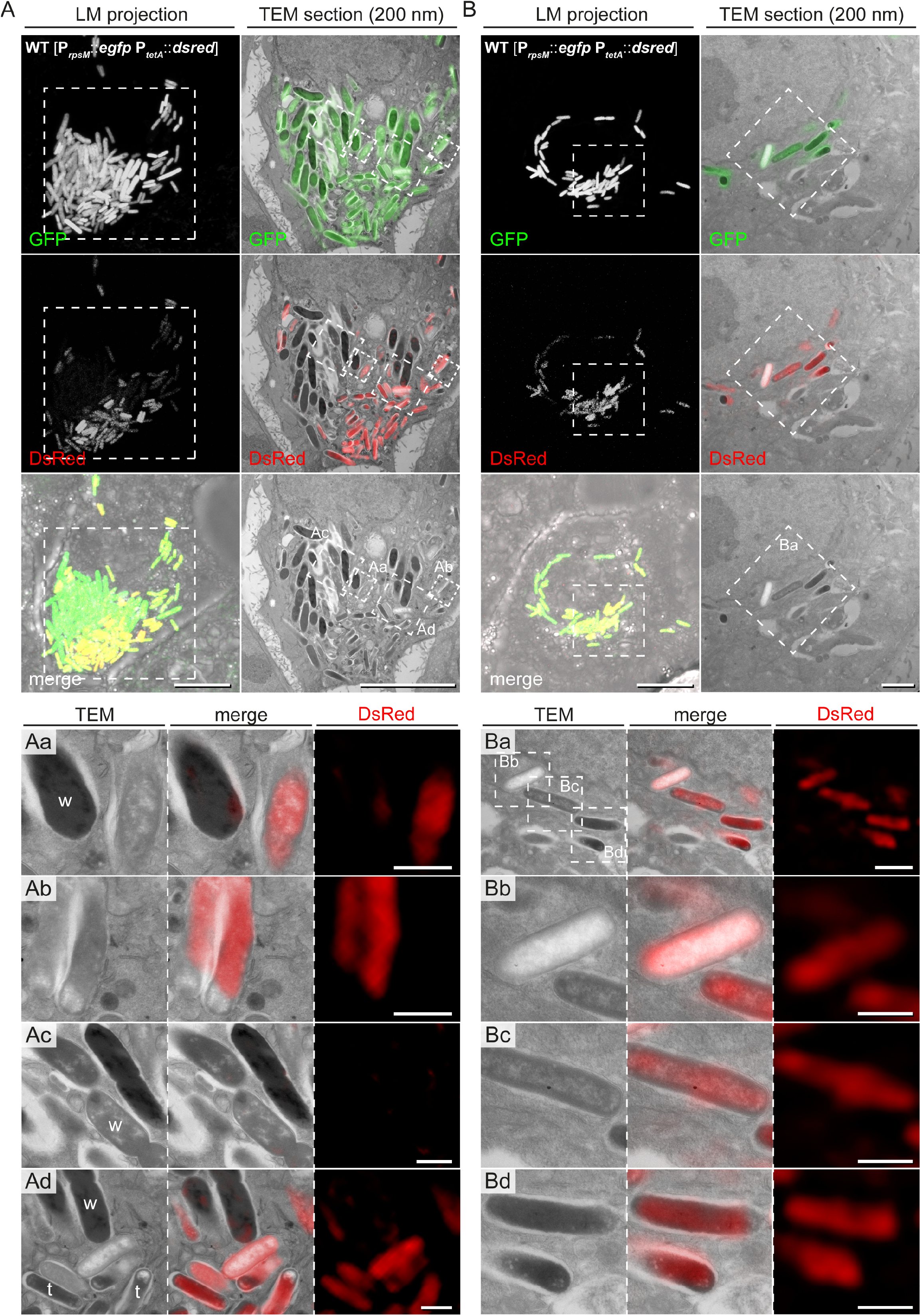
CLEM reveals ultrastructural and metabolic diversities of hyper-replicating intracellular STM WT in HeLa cells. HeLa cells seeded on gridded cover slip were infected with STM WT harboring the dual-color vitality reporter and visualized 16 h p.i. *Gfp* was constitutively expressed in STM WT while *dsred* expression was induced by AHT 2 h prior to imaging by confocal FM. Subsequently, cells were processed for TEM. **A**) Representative HeLa cell with hyper-replicating intracellular STM WT, either DsRed-positive (active cells) or DsRed-negative (inactive). **B**) Representative HeLa cell with replicating STM WT, which are DsRed-positive. For CLEM, EF-TEM micrographs of 200 nm thick sections of infected HeLa cells obtained at 120 keV were correlated with GFP or DsRed fluorescence signals of confocal sections after deconvolution. **a-d**) High-resolution CLEM of ROIs (white boxes) harboring intracellular STM WT of diverse electron density (TEM) and activity levels (DsRed, FM). Note ED and EL STM during division (**Ac**), which are DsRed-negative. ‘Wide’ (w) and ‘thin’ (t) STM are marked. Scale bars, 10 μm (**A**, all overviews and **B**, LM overviews), 1 μm (**B**, TEM overviews, **Aa-Ad** and **Ba**), 500 nm (**Bb-d**).

All together, these data provide further evidence for the existence of different ultrastructural classes. Furthermore, we found STM cells with special features (Figure 7). It had clear condensations of structures in the cytoplasm with a dense layer surrounding loose materials in the center (halo-like condensation). Correlation of fluorescence signals with the ultrastructural profile showed that the ‘halo’-like electron density contained GFP and DsRed. The biosynthetic activity of this cell was at high level, suggesting high vitality. At the poles, lucent blebs of regular size and shape were visible suggesting that they were not lysis spots. The inner bacterial membrane in proximity of lucent blebs was intact (Figure 7BC).

**Figure 7:**
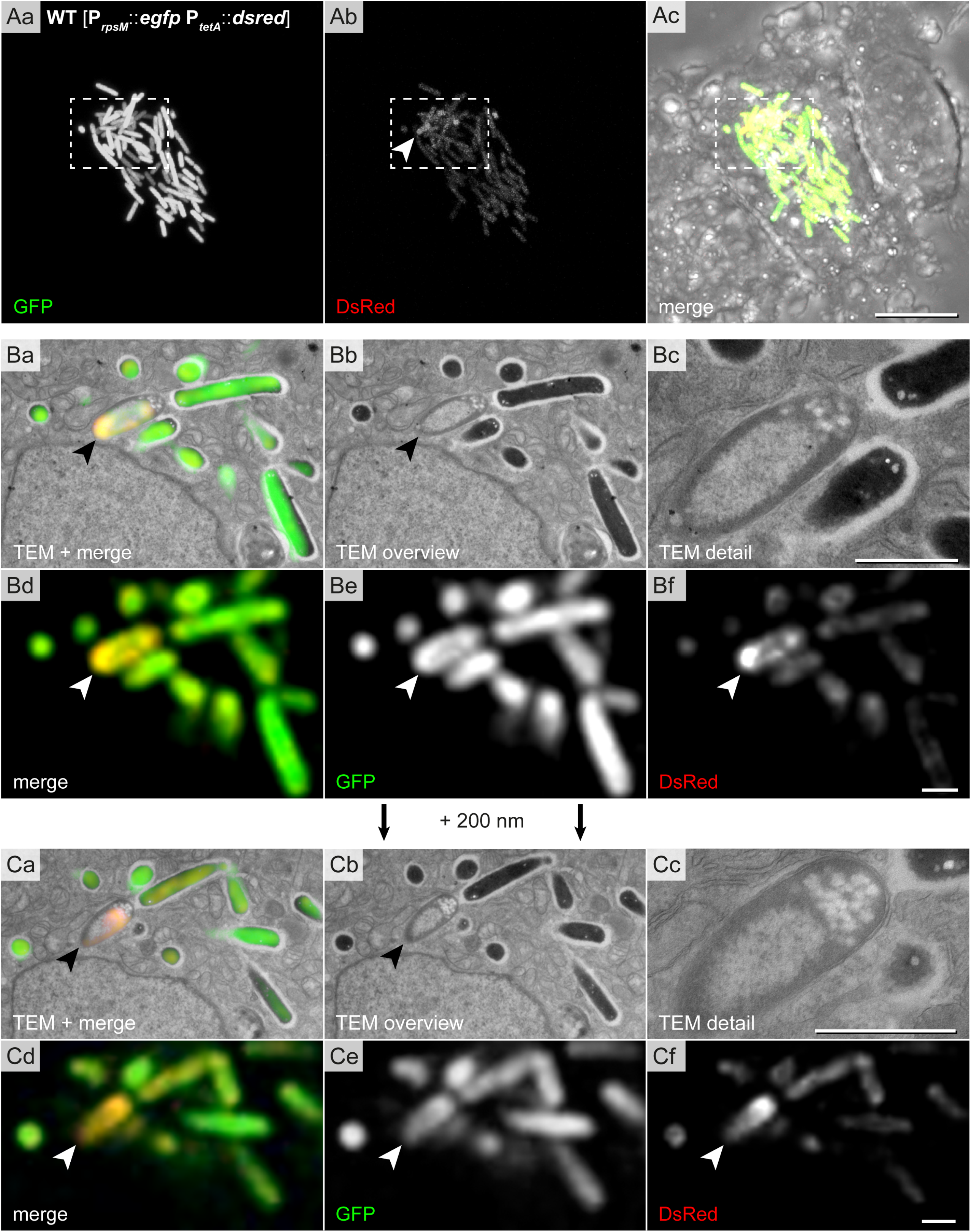
High-resolution CLEM of intracellular STM WT precisely locates proteins in bacterial cytoplasm: halo-like distribution of GFP/DsRed and electron density in single cells. Intracellular STM WT harboring dual-color vitality reporter was visualized inside HeLa cells seeded on gridded cover slips 16 h p.i. STM WT constitutively expressed *gfp*, while *dsred* expression was induced by AHT 2 h prior to imaging by confocal FM. Subsequently, cells were processed for TEM. **Aa-Ac**) Overview of HeLa cell with hyper-replicating intracellular STM WT (GFP, green) showing metabolic activity (DsRed, yellow in merge). Dashed boxes indicate CLEM region in **B** and **C**. **B, C**) CLEM of consecutive 200 nm thick sections of region with STM WT of halo-like electron density (indicated by arrowheads). GFP and DsRed confocal fluorescence signals after deconvolution (**d-f**) are correlated with 120 keV EF-TEM micrographs. GFP and DsRed are distributed in a halo-shaped electron density. Scale bars, 10 μm (**A**), 1 μm (**B, C**).

Hence, the CLEM approach had sufficient resolution to visualize distinct DsRed/GFP distributions inside bacteria in correlation with the ultrastructure.

### The ‘halo’ type of bacteria

We noticed presence of STM with halo-like condensations in the cytoplasm also in o/n LB cultures, what allowed for quantifications of this type. Cells with halo-like condensation had centrally located lucent region, occupying 39-55% of the whole cell area (48.5% ± 4.9). Quantification revealed that cultures in late-logarithmically growth did not contain any cells with halo-like condensations, in contrast to stationary cultures (o/n) (Figure 8A). Moreover, the Δ*sodAB* strain more frequently formed halo-like structures, suggesting a link between stress and ‘halo’ ultrastructure (Figure 8A). In contrast, we did not detect halo-like condensations in any STM cell cultured in PCN medium, independently of growth phase, pH, AA supplementation, or even shock conditions.

**Figure 8:**
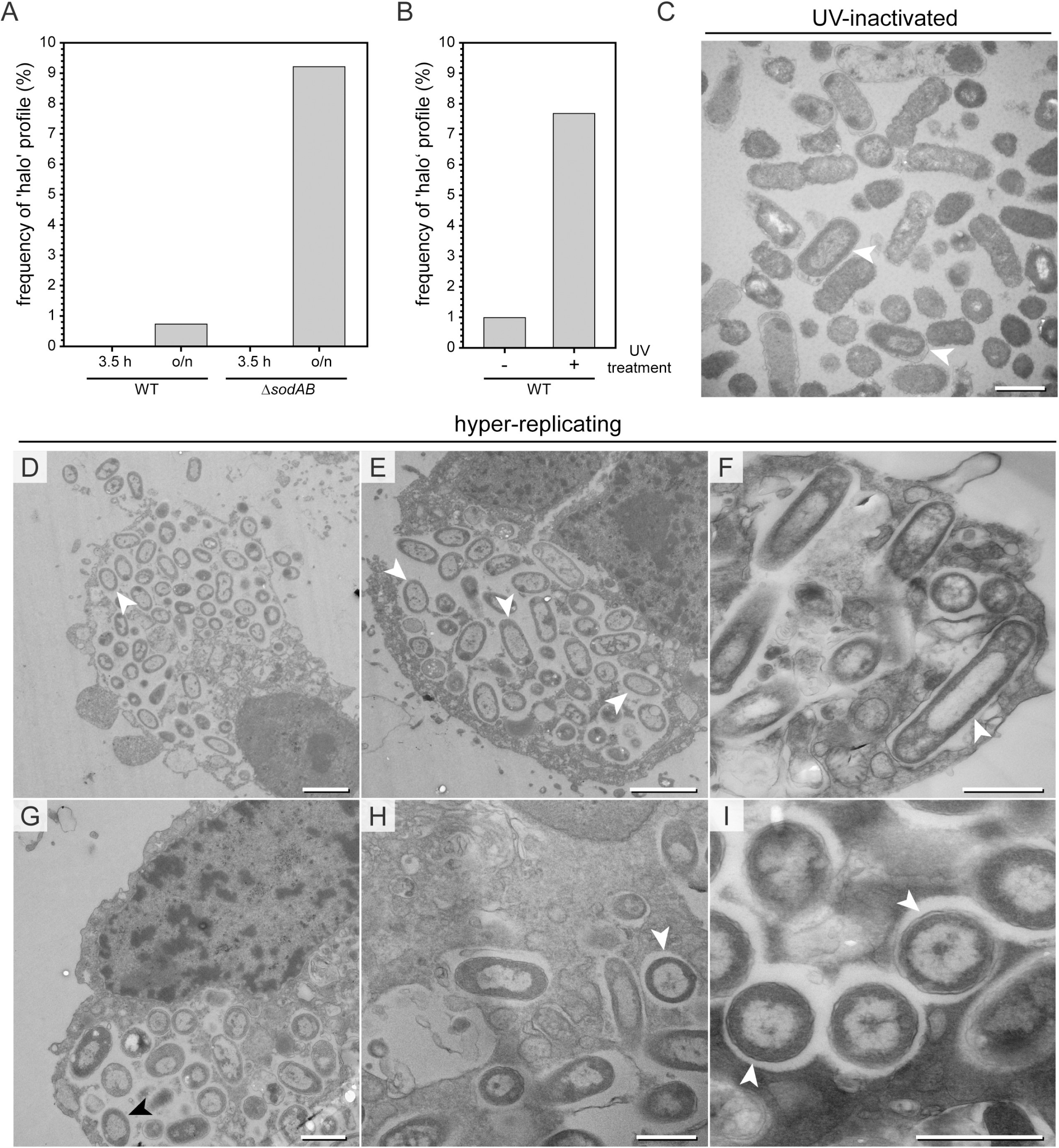
‘Halo’ type of STM WT dominates in critical environmental conditions. **A**) Comparison of relative numbers of halo-shaped STM WT and Δ*sodAB* cultured in LB medium overnight (o/n) or further subcultured for 3.5 h in fresh medium prior to TEM preparation (TEM in Fig. S1D-G). Number of quantified cells: 1,032, 819, 747, and 510, for WT 3.5 h, WT o/n, Δs*odAB* 3.5 h, and Δ*sodAB* o/n, respectively. **B, C**) STM WT was irradiated 60 sec with UV light (305 nm) prior to TEM preparation. **B**) Comparison of relative numbers of halo-shaped STM WT after UV irradiation (arrowheads in **C**) to its control, the same LB culture without UV treatment. Numbers of quantified cells: 716 and 612, for UV-treated and non-treated groups, respectively. **C**) TEM micrograph of UV-treated STM WT with halo-shaped profiles (indicated by arrowheads). **D-I**) TEM micrographs of HeLa cells 16 h p.i. with STM WT. Infected HeLa cells containing hyper-replicating STM WT show disrupted cell membranes and signs of cell death. Nearly all intracellular STM WT have cytoplasmic densities distributed as a halo (arrowheads). Scale bars, 2.5 μm (**D, E**), 1 μm (**C, F-I**).

We further tested to induce experimentally the appearance of the halo-like type. UV light of wavelengths of 290-320 nm (UVB), or 254 nm (UVC) are highly bactericidal (54). Therefore, STM WT cultured in LB medium was analyzed by TEM after UV treatment, which resulted in complete lack of colony growth on agar plates (Figure 8B-C and S6A-E). The ultrastructural profiles of UV-treated STM were classified, revealing reduced numbers of EL type when compared to untreated cells. ED profiles were slightly more frequent, together with profiles showing signs of cellular death. UV treatment induced formation of the halo-like type to almost the frequency observed for STM Δ*sodAB* (Figure 8ABC, arrowhead). Hence, there is a correlation of the halo-like type frequency with occurrence of cell death induced by UV illumination.

We also determined biosynthetic capacity after UV treatment and found significant reduction in DsRed production (Figure S6E). Surprisingly, DsRed was synthesized after UV treatment, while no colony growth was observed (Figure S6EF). We analyzed infected host cells with respect to presence of halo-like morphotype and found that this type dominated intracellular populations in case of hyper-replication, when bacteria were leaving the host, or host cells were necrotic (Figure 8D-I, arrowheads). In infection experiments, the antibiotic gentamicin is used to kill extracellular bacteria, while intracellular bacteria are protected. We tested if gentamicin could be a cause for the increase of the halo-like type in infection experiments, however, treatment of STM cultures with gentamicin did not recapitulate this result (data not shown).

After growth in rich media like LB cultures, halo-like morphotype was about 10%, suggesting low exposure to cell-damaging agents. Within host cells, the impact on ultrastructural characteristics of STM was more dramatic. Taken together, these results demonstrate that the halo-like type could be an indicator of exposure of STM to critical environmental conditions.

## Discussion

In this work, we present evidence that several morphotypes of viable bacteria exist. Based on ultrastructural criteria, these morphotypes differ significantly in cytoplasmic electron density, which is well distinguishable at low EM resolution (1,000-2,000-fold magnification, 50 keV), as well as in organization and visibility of defined (chromosome, ribosomes) and undefined structures (dense cytoplasmic areas). We present EL and ED types of STM, representing viable bacteria active for protein biosynthesis and/or able to divide. The frequency of morphotypes in a bacterial population can be used as an indicator of environmental changes, what in turn enables new ways of EM data interpretation. Moreover, ultrastructural heterogeneity is not limited to *Salmonella enterica*, since published TEM micrographs of other bacteria demonstrate morphotypes similar to EL, ED, as well as halo-like type. These types seem to be neither restricted to Gram-negative nor pathogenic bacteria. Their examples have been visualized intracellular in both, cell cultures and infected animals previously, and were not restricted to mammalian cells (22–34). However, the nature or function of the ultrastructural diversity remained enigmatic.

Here, we provide evidence that the ultrastructural heterogeneity, other than death-related, is associated with environmental changes. We further demonstrated that the external AA supplementation is required for ED induction and that the oxidative stress-dependent formation of ED bacteria only takes place in nutrient-rich medium. In contrast, ED cells were hardly inducible by oxidative stress in minimal media, which did not contain AA. Furthermore, induction of ED type is pH sensitive with restriction to a phagosomal pH. Consistently with weak survival capabilities, the lack of obvious ED type in PCN medium despite oxidative stress, may reflect difficulties of cells to respond properly. Recently, a morphotype reminiscent with ED has been described in *E. coli*, as representative of healthy cells (12, 55). In addition, we have identified the very characteristic halo-like type, which dominates bacterial populations exposed to extreme conditions. This type occurred sporadically in stationary phase and is increased in cultures after bactericidal UV treatment, pointing to its link with deadly noxes. CLEM analysis of halo-like type suggests that the dense matter around the DNA-containing center is of proteinaceous nature since GFP and DsRed were located within this area. At this moment we do not know the underlying mechanisms of different cytoplasmic states at the ultrastructural level but one possible explanation for ED type can be macromolecular crowding as a result of stress response (56, 57). Previously, molecular crowding has been demonstrated in *E. coli* after osmotic up-shift, affecting diffusion of GFP. These changes were accompanied by dramatic cytoplasmic shrinkage, which was only sporadically observed at the cell poles by TEM in our experiments (58, 59).

Global and dramatic ultrastructural alterations of bacteria occurred very fast, because differences were visible by TEM when cells were fixed directly after stress treatment. On the other side, dead and dying bacterial cells, according to ultrastructural criteria like membrane injuries, lysis, and/or leakage of content are not frequent, even after bactericidal UV applications or after shock conditions. This suggests that STM can be alive for a long time without being capable to form colonies. We also visualized cells at high resolution to clearly discriminate between lysis and storage granules or other electron-lucent components not related with death (this study; 19, 58), and between molecular condensation and protein-based organelles like carboxisomes (19, 60). Except ruptures and detachments, membrane waving and surface protrusions cannot be interpreted with convenience as indicators of deadly membrane stress, since they may reflect adaptive activity of cells, i.e. changes in membrane fluidity or vesicle formation (61–66). Criteria for dying cells were found in EL, ED and halo-like type, exhibiting severity stages as intermediates of death processes, different cellular location and mix of criteria.

Taken together, our study sheds more light on ultrastructural heterogeneity of STM and revealed possible EM indicators, which allow to broaden EM data interpretation. Since the ED type could serve as indicator for oxidative stress, and the halo-like type as an indicator for hazardous environmental conditions, future research infection in biology will benefit, because these indicators may be used when detailed analyses are difficult or impossible. For example, when bacterial populations *in vivo* in deep tissues are investigated or populations not compatible with reporter systems are analyzed as for example free-living microorganisms, which were directly analyzed without cultivation.

## Acknowledgements

This work was supported by the DFG by grant P15 and the Z project in SFB 944. The excellent technical assistance of Birgit Hemmis and Britta Brickwedde is kindly acknowledged.

## Materials and Methods

### Bacterial strains and growth conditions

*Salmonella enterica* serovar Typhimurium strain NCTC12023 (STM) was used as wild-type strain and isogenic strain MvP2400 (Δ*sodA*::FRT Δ*sodB*::FRT) has been described (41). Bacteria were cultured in lysogeny broth (LB), or PCN medium supplemented with 0.4 mM (for paraquat treatment) or 25 mM PO_4_^−^ (44, 45) at pH of 5.8 or 7.4 at 37 °C with aeration. Optionally, medium was supplemented with an 1 x mix of 20 amino acids (alanine (0.8 mM), arginine (5.2 mM), asparagine (0.4 mM), aspartate (0.4 mM), cysteine (0.1 mM), glutamic acid (0.6 mM), glutamine (0.6 mM), glycine (0.8 mM), histidine (0.2 mM), isoleucine (0.4 mM), leucine (0.8 mM), lysine (0.4 mM), methionine (0.2 mM), phenylalanine (0.4 mM), proline (0.4 mM), serine (10.0 mM), threonine (0.4 mM), tryptophan (0.1 mM), tyrosine (0.2 mM), valine (0.6 mM)) (44). When required, carbenicillin or chloramphenicol were added at 50 μg x ml^−1^ or 200 μg x ml^−1^, respectively. For live cell imaging and flow cytometry analysis of bacterial metabolic activity, the strains harbored plasmid pWRG658 (P_*rpsM*_::*gfpmut3A tetR* P_*tetA*_::*dsRed T3_*S4T) (51) for constitutive expression of *gfp* and AHT-inducible expression of *dsred*.

### AHT induction

AHT (Fluka, Sigma-Aldrich) stock solutions were stored in aliquots of 200 μg x ml^−1^ in dimethylformamide (DMF) at −20 °C in the dark. For induction of expression of the P_*tetA*_-controlled dual-color vitality sensor, AHT was added directly to LB broth or cell culture medium to a concentration of 100 ng x ml^−1^ as indicated.

### Propidium iodide staining of STM

Propidium iodide (PI) (Sigma-Aldrich) was used as described (67) to analyze cell envelope integrity at a concentration of 30 μM in PBS. STM was cultured as indicated, PI was added, and incubated for 10 min in the dark. Subsequently, bacteria were washed twice by centrifugation (5,000 x g, 5 min) in the same buffer, 5 μl of diluted bacteria were put on a glass slide, covered with a cover slip and imaged by the Zeiss LSM (Zeiss).

### Stress induction by methyl viologen, heat, hyper- or hypo-osmolarity

STM strains were cultured overnight as indicated, diluted 1:31 in fresh medium, subcultured for further 3.5 h, shifted to fresh PCN medium as indicated and exposed to methyl viologen (Sigma-Aldrich) for 1 h at RT without shaking, to 80 °C (in PCN medium pH 7.4) or to hyper-osmolar (PCN medium pH 7.4 containing 600 mM NaCl) or hypo-osmolar conditions (pure H_2_O_dd_) for 2 h. Effect of methyl viologen was confirmed by plating of bacteria onto LB plates. Subsequently, bacteria were processed for PI staining or TEM as described above or below, respectively.

### UV inactivation of STM

Overnight cultures of STM were grown in LB broth, normalized to an OD_600_ of 0.2 in PBS and transferred to a petri dish following irradiation with UV light (305 nm) for 60 sec. Successful UV inactivation was always confirmed by plating of inactivated bacterial suspension onto LB plates. If indicated, inactivated bacteria were processed for EM as described below.

### Flow cytometry analysis

Overnight cultures of STM were grown in LB broth, diluted 1:31 in fresh LB and subcultured for further 6 h. At indicated time points, samples were taken, diluted in PBS and directly subjected to flow cytometry without fixation. Flow cytometry was performed on an Attune NxT instrument (ThermoFischer Scientific) at a flow rate of 25 μl x min^−1^. At least 50,000 bacteria were gated by virtue of the constitutive GFP fluorescence. The percentage of DsRed-positive bacteria was determined, the intensity of the DsRed fluorescence per gated STM cell was recorded and x-medians for DsRed intensities were calculated.

### Cell lines and cell culture

For infection experiments the non-polarized epithelial cell line HeLa (American Type Culture Collection, ATCC no. CCL-2) stably transfected with LAMP1-GFP was used. HeLa cells were cultured in Dulbecco’s modified Eagle’s medium (DMEM) containing 4.5 g x l^−1^ glucose, 4 mM stable glutamine and sodium pyruvate (Biochrom) and supplemented with 10% inactivated fetal calf serum (iFCS) (Sigma-Aldrich) at 37 °C, 5% CO_2_ and 90% humidity.

### Host cell infection

For infection of HeLa LAMP1-GFP cells, *Salmonella* strains were grown overnight in LB broth, diluted 1:31 in fresh LB and subcultured for further 3.5 h to induce maximal invasiveness. Infection was performed with a multiplicity of infection (MOI) of 50 for 25 min at 37 °C, 5% CO_2_ and 90% humidity. Subsequently, cells were washed thrice with PBS and incubated for 1 h with medium containing 100 mg x ml^−1^ gentamicin (Applichem) to kill all non-invaded bacteria. Afterwards, the medium was replaced by medium containing 10 mg x mL^−1^ gentamicin until the end of the experiment.

### Live cell imaging and image deconvolution

For live cell imaging, DMEM was replaced by imaging medium consisting of Minimal Essential Medium (MEM) with Earle’s salts, without NaHCO_3_, without L-glutamine and without phenol red (Biochrom) supplemented with 30 mM HEPES (4-(2-hydroxyethyl)-1-piperazineethanesulfonic acid) (Sigma-Aldrich) with a pH of 7.4. For imaging of fixed cells, cells were washed thrice with PBS and incubated for 15 min with PBS containing 3% *para*-formaldehyde (PFA) to ensure complete fixation of cells. Subsequently, cells were washed thrice with PBS and blocked with blocking solution containing 2% bovine serum albumin and 2% goat serum in PBS. Fluorescence imaging was performed using the confocal laser-scanning microscope (CLSM) Leica SP5. For setting adjustment, image acquisition and image processing the software LAS AF (Leica, Wetzlar, Germany) was used. Image acquisition was performed using objectives 10x (HC PL FL 10x, NA 0.3), 20x (HC PL APO CS 20x, NA 0.7), 40x (HCX PL APO CS 40x, NA 1.25–0.75) and 100x objective (HCX PL APO CS 100x, NA 1.4–0.7) (Leica, Wetzlar, Germany) and the polychromic mirror TD 488/543/633 for the three channels GFP, DsRed and DIC. For CLEM experiments, images were further deconvoluted using Huygens software (Scientific Volume Imaging B.V., Hilversum, The Netherlands) to better correlate the expression patterns of DsRed to the bacterial ultrastructure. Live cell imaging was performed using the Zeiss Cell Observer microscope with Yokogawa Spinning Disc Unit CSU-X1a, Evolve EMCCD camera (Photometrics, USA) and live cell periphery, equipped with an Alpha Plan-Apochromat 63x (NA 1.46) oil immersion objective (Zeiss, Oberkochen, Germany). Following filter combinations were used for image acquisition: GFP with BP 525/50, DsRed with LP 580 and processed by the ZEN 2012 (Zeiss, Oberkochen, Germany) software. Scale bars for all acquired images were added with Photoshop CS6 (Adobe).

### Sample preparation for TEM

STM cultured either in LB or PCN medium were fixed with 2.5% glutaraldehyde (GA) (Electron Microscopy Science) in 100 mM phosphate buffer (81.8 mM Na_2_HPO_4_ and 18.2 mM KH_2_PO_4_, pH 7.2) over night at 4 °C. Unreacted aldehydes were blocked with 100 mM glycine in buffer for 15 min. Osmification was performed with 1% osmium tetroxide (Electron Microscopy Science) in 100 mM phosphate buffer for 60 min on ice following washing several times with phosphate buffer and ultrapure water (MilliQ). Subsequently, contrasting with 1% uranyl acetate (Electron Microscopy Science) in MilliQ for 30 min was performed following several washing steps. Afterwards, cells were dehydrated in a cold graded ethanol series finally rinsing once in anhydrous ethanol and twice in anhydrous acetone at room temperature. Infiltration was performed in mixes of acetone and EPON812 (Serva). After every incubation or washing step bacteria were centrifuged (2000 x g, 3 min), the supernatant was discarded followed by the next preparation step.

### Sample preparation for CLEM

Two days prior to infection HeLa LAMP1-GFP cells (1 × 10^5^) were seeded onto a gridded coverslip in a petri dish (MatTek, Ashland, MA). 14 h p.i. 100 ng x ml^−1^ AHT was added to the cells for induction of reporter plasmid. 16 h p.i. cells were pre-fixed with pre-warmed 2.5% GA in 0.1 M phosphate buffer for 15 min at 37 °C. After washing the cells thrice with PBS, ROIs were documented and images were acquired. Subsequently, further fixation was performed using 2.5% GA in 0.1 M phosphate buffer over night at 4 °C. Quenching, osmification and contrasting was performed as described above. Then, the gridded coverslip was removed from the petri dish and was transferred to a glass dish. Afterwards, cells were dehydrated in a cold graded ethanol series, finally rinsing once in anhydrous ethanol and twice in anhydrous acetone at room temperature. Infiltration and flat-embedding were performed in mixes of acetone and EPON812 (Serva). During the removal of the gridded coverslip from the polymerized EPON the engraved coordinates were transferred to the EPON surface and allowed easy relocation by microscopy. ROIs were cut using a scalpel and were transferred to an EPON block. Serial 200 nm sections were generated by an ultramicrotome (Leica EM UC7) and collected on formvar-coated copper EM slot grids.

### Transmission electron microscopy

High-resolution analysis including CLEM was performed using the Libra 120 TEM (Zeiss, Oberkochen, Germany) operating at 120 keV and equipped with an Omega energy filter and a 2,000×2,000-pixel digital camera (Troendle). In addition, TEM was performed using a Zeiss 902 system (Zeiss, Oberkochen, Germany) operating at 50 keV. Images were taken with the software ImageSP (TRS image SysProg, Moorenwies, Germany). TEM micrographs were adjusted for brightness and contrast enhanced using ImageJ or Photoshop software when necessary. For image analysis, software ImageJ (http://rsbweb.nih.gov/ij/) was used. Stitching and overlay of CLSM and TEM images were done using Photoshop CS6 (Adobe).

## Suppl. Figure Legends

**Figure S1 (related to Fig. 1): A-C**) Quantification of data obtained by 120 keV EF-TEM imaging (related with Fig. 1C-E). **A**) Comparison of electron densities (difference values of MGV of STM cytoplasm to MGV of background) of ED and EL STM WT in different regions of sample. Averages of electron densities (**B**), and **(C**) averaged density ratio (mean ± SD). **D-M**) Electron micrographs at 50 keV of STM WT and Δ*sodAB* strains from o/n cultures or 3.5 h subcultures in LB medium (**D-G**), STM WT from o/n or 3.5 h subcultures in PCN, pH 7.4 with (**J, K**), or without AA supplementation (**H, I**), or in PCN, pH 5.8 (**L, M**). Arrows indicate cells with mixed electron densities and halo-like distribution in **E**. Arrowheads in **G** indicate cells of STM Δ*sodAB* with translucent spot not found in 3.5 h subculture of the same culture. Scale bar, 1 μm. Statistical analysis was accomplished by Student’s *t*-test, and significance levels are indicated as follows: *, p < 0.05; **, p < 0.01; ***, p < 0.001; n.s., not significant.

**Figure S2 (related to Fig. 2): PQ treatments affects membrane integrity to similar extend in STM WT and Δ*sodAB* strains**. **A**) Relative numbers of PI-positive STM Δ*sodAB* without and after treatment with PQ. Obvious increase of PI-load events in culture was achieved after treatment with 50 μM PQ in comparison to control (no treatment) and treatment with PQ at concentrations of 5 μM or 10 μM. Number of cells quantified: 3,876, 1,5825, 15,804, and 18,016, for 0 μM, 5 μM, 10 μM, and 50 μM PQ, respectively. **B, C**) Comparison of PQ effects at 50 μM PQ between STM WT and Δ*sodAB*: the relative number of PI-positive cells and fluorescence intensities of PI. PI was loaded 12 h after PQ treatment. STM Δ*sodAB* shows high variability in number and degree (intensities) of PI-load events in the culture (controls). Statistical analysis was accomplished by Student’s *t*-test and significance levels are indicated as follows: *, p < 0.05; **, p < 0.01; ***, p < 0.001; n.s., not significant.

**Figure S3 (related to Fig. 3): Treatments of STM WT using various stressors have impact on ultrastructure. A-F**) Electron micrograph of STM WT cultured in PCN, pH 7.4 for 3.5 h and shifted to fresh PCN, pH 7.4 (**A-C**), or PCN, pH 5.8 (**D-F**) for incubation without PQ (control **A, D**), with 100 μM PQ (**B, D**), or 500 μM PQ (**C, F**). **G-L**) EF-TEM micrographs of STM WT without (control), after 100 μM or 500 μM PQ treatment, displayed as an electron density scale. Arrowheads point to cells with denser cytoplasm compared to controls. **M-O**) Electron micrograph of STM WT cultured in PCN, pH 7.4 supplemented with AA for 3.5 h and shifted to fresh PCN, pH 7.4 for incubation without (control **G**), with 100 μM PQ (**H**), or 500 μM PQ (I). **P-S**) Electron micrograph of STM WT cultured in PCN, pH 7.4 or PCN, pH 5.8 supplemented with AA for 3.5 h and shifted to fresh PCN, pH 3.0 for incubation without (**P** and R), or with 1 mM PQ (**Q** and **S**). **T-V**) Electron micrographs of STM WT cultured in PCN, pH 7.4 for 3.5 h and shifted to fresh PCN, pH 7.4 for incubation at 80 °C (**T**), to PCN, pH 7.4 containing 600 mM NaCl (**U**), or to pure H_2_O_dd_ (**V**). **W**) Aliquots of STM subcultured for 3.5 h in PCN, pH 7.4 or PCN, pH 5.8 without PQ treatment, or treatment with 100 μM, 500 μM or 1 mM PQ were plated onto agar plates, and CFU were determined. CFU per ml culture are expressed as percentage of CFU of untreated culture. **X, Y**) Relative numbers of STM WT with shrinkage (arrowhead in **D**) or features of lysis (arrowhead in **F**) in indicated culture conditions. Features of untreated control samples in % of total are shown in **X** and x-fold increase of features of stressor-treated samples compared to the respective untreated sample is shown in **Y**. In addition, values of shrinkage and lysis of stress-treated samples in % of the total population is indicated above. Quantified cells for each condition: 100-300 cells. Scale bars, 1 μm (**A-V**), 250 nm (detail in C-F).

**Figure S4 (related to Fig. 5–7): Functional characterization of dual-color vitality reporter of STM WT. A**) The dual-color vitality reporter harbors a constitutive *gfp* expression and an AHT-inducible expression of *dsred*. Metabolically active bacteria are expected to synthesize DsRed after the addition of AHT, in contrast to metabolically inactive bacteria, or when protein biosynthesis is experimentally blocked by chloramphenicol (Cm) after induction. **B**) Cytometric gating shows detection of bacteria-sized particles selected by FSC/SSC. GFP-positive cells were gated, the DsRed fluorescence intensity of the GFP-positive bacterial population was determined, and the x-median RFI of the entire population was calculated. **C**) Comparison of relative DsRed-positive numbers of STM WT subcultures without AHT induction, with AHT induction or AHT induction/Cm inhibition. DsRed-positive counts achieve a maximum already 1 h after addition of AHT. Cm inhibits synthesis of DsRed after AHT induction. The x-median represents the AHT-induced DsRed signal of the GFP-positive bacterial population. **D**) DsRed intensity peaks at 0 h (black), 3 h (red, start of AHT induction), or 7 h (green) of subculture. DsRed intensity shifts towards higher values after addition of AHT (upper histograms). If no AHT is added (middle histograms), or protein biosynthesis is blocked by addition of Cm (lower histograms), DsRed intensity remains low. **E**) Intracellular STM WT with dual-color vitality reporter (GFP) in infected HeLa cells were DsRed positive (yellow in merge) only after addition of AHT. Addition of Cm with AHT successfully blocks DsRed synthesis. Scale bar, 5 μm.

**Figure S5 (related with Fig. 6 and 7): CLEM approach to analyze intracellular STM at high resolution. A**) To register coordinates for CLEM, HeLa cells were seeded at a gridded cover slip prior to infection with STM WT harboring the dual-color vitality reporter. *Gfp* was constitutively expressed in STM WT, while *dsred* expression was induced by addition of AHT 2 h prior to imaging of selected ROIs by CLSM at 16 p.i. Subsequently, samples were fixed and prepared for TEM by flat-embedding in plastic resin. Coordinates from CLSM were well visible, allowing relocation of ROIs. Resin fragments containing ROIs were dissected and individually fixed to resin blocks for serial 200 nm ultra-sectioning. After relocation of HeLa cells containing STM WT in TEM modality, serial sections were acquired at 120 keV with energy filtering (Ω filter). This approach significantly reduced time needed for sample preparation to image whole cells with TEM, providing the opportunity to collect data of high quality at optimal resolutions for correlation or TEM analysis. **B**) CLEM imaging steps: visualization by CLSM of intracellular STM WT by virtue of GFP fluorescence, and evaluation of metabolic activity by virtue of DsRed intensity (Z-stack through STM population, maximum intensity projection is shown), registration of positions of infected HeLa cells using BF-LM, relocalization of HeLa cells in resin using imprinted CLEM coordinates, TEM imaging, correlation of single CLSM planes with corresponding single TEM images. **C**) Comparison of fluorescence signals without and with deconvolution using Huygens software. After deconvolution, location of proteins GFP and DsRed was increased, and signal-to-noise ratio was improved. Scale bars, 20 μm (**B**, left panel), 5 μm (**B**, right panel and **C**), 2 μm (**C**, detail).

**Figure S6 (related with Fig. 8): UV treatment of STM WT boosts ultrastructural diversity. A, B**) Cultures of STM WT grown o/n in LB were irradiated 60 sec. with UV light (305 nm) prior preparation for TEM. As control, STM WT of the same culture was left without irradiation and processed in parallel for TEM. Various ultrastructural profiles were defined: (**I**) EL, (**II**) ED, (**III**) initiated cell death, indicated by shrinkage and damage of inner membrane, or (**IV**) presence of electron-dense (condensation) and translucent (lysis) spots in the bacterial cytosol, (**V**) dead cells with condense cytosolic materials and severally fragmented or lacking inner membranes, and (**VI**) the halo-shaped profile with electron densities at peripheries of cytoplasm. **C**) Comparison of relative numbers of profiles **I** to **V** of UV-irradiated and untreated STM WT. **VI**-type relative number is shown in Fig. 8B. Number of quantified cells: 716 and 612 for UV-irradiated and non-treated groups, respectively. **D**) Comparison of accumulated frequency of STM with **I** and **II** profiles (ED+EL), to accumulated frequency of STM with morphological impairments (**III**-**V**, shrinkage and lysis of various severity). **E**) Effect of UV irradiation on bacterial survival. Plating of aliquots of STM cultures onto agar plates and CFU determination was performed for all experiments involving UV irradiation. **F**) Flow cytometry analysis of STM WT activity using dual-color vitality reporter after UV irradiation. Overnight cultures of STM WT were UV-treated or untreated (control) prior to subculture in fresh LB medium. Induction of *dsred* expression was initiated by addition of AHT at start of subculture. Cultures were sampled in hourly intervals with parallel plating onto agar plates for CFU counts. The x-median RFI represents the AHT-induced DsRed signal of the GFP-positive bacterial population. **G**) Viability of STM analyzed in **F**. The amount of living bacteria was calculated and normalized to 100% of living STM in an untreated sample. As control, STM without UV irradiation were used. Note an increase of DsRed intensity of UV-irradiated bacteria (**F**) in LB medium although their growth on LB plates is totally inhibited (**G**). Scale bars, 5 μm (**A, B**), 500 nm (**I-VI**).

